# Natural selections on both amino acid sequences and expression levels are determinants of ohnolog retention

**DOI:** 10.1101/2021.02.24.432806

**Authors:** Lin Miao, Miaoxin Li

## Abstract

The mechanism of ohnolog retention is a subject of concern in evolutionary biology. Natural selections on coding sequences and gene dosages have been proposed to be determinants of ohnolog retention. However, the relationship between the two models is not widely accepted, and the role of regulatory sequences on ohnolog retention has long been neglected. In this study, based on a model of complex traits’ genetic architecture, we compared the natural selection’s strength on corresponding sequences between ohnologs and non-ohnologs by comparing complex traits’ heritability enrichments. We showed that complex traits’ regulatory sequences’ heritability enrichments (*p* = 1.1 × 10^−5^ in 5 kb flanking regions) and expression-mediated heritability enrichments (*p* = 2.1 × 10^−5^) of ohnologs were significantly higher than non-ohnologs. Then, we deduced that regulatory sequences of ohnologs were under substantial natural selection, which was also a determent of ohnolog retention. Meanwhile, we showed that in coding sequences, the complex traits’ heritability enrichments of ohnologs were significantly higher than of non-ohnologs (*p* = 9.9 × 10^−5^), supporting the ohnolog retention model of natural selection on coding sequences. We also showed that complex traits’ causal gene expression effect sizes of ohnologs were significantly larger than of non-ohnologs (*p* = 8.8 × 10^−6^), supporting the ohnolog retention model of natural selection on gene dosages. In conclusion, we provide the first unified framework to show that both amino acid sequences and expression levels of ohnologs are under substantial selection, which may end the long-standing debate on ohnolog retention models.

## Introduction

Ohnologs are paralogs that descended from two rounds of whole-genome duplications (WGD) early in the evolutionary history of Vertebrata. The types of natural selection that ohnologs have undergone are closely linked to the types of pathogenic variants observed in ohnologs, which makes the role of ohnologs in disease and the mechanism of ohnolog retention a subject of concern. Due to the observation of overrepresented ohnologs in pathogenic copy number mutations, ohnologs were considered dosage-sensitive, and natural selection on gene dosages was proposed to be a driving force behind the retention of ohnologs (***Makino and McLysaght, 2010; McLysaght et al., 2014; Rice and McLysaght, 2017***). Meanwhile, because human dominant disease genes were enriched in ohnologs, ohnologs were considered to be prone to dominant deleterious mutations, and purifying selection on coding sequencings was thought to be the driving force behind the retention of ohnologs (***Singh et al., 2012, 2014***; ***Roux et al., 2017***). However, the relationship between the two models of ohnolog retention is unclear, and the role of ohnolog regulatory sequences in human complex traits and regulatory sequences’ role on ohnolog retention has long been neglected.

In this study, we take advantage of Eyre-Walker’s model, in which mutations that affect a complex trait also affect fitness (***Eyre-Walker, 2010; Keightley and Hill, 1990***), to deduce the relative strength of natural selection on corresponding sequences between ohnologs and non-ohnologs. We tested the null hypotheses that there were no differences between complex traits’ heritability enrichments of corresponding sequences (e.g., intron) between ohnologs and non-ohnologs. We deduced if the natural selections on corresponding sequences (e.g., intron) of ohnologs were stronger than non-ohnologs based on the results of hypothesis tests.

## Results

### Comparison of complex traits’ heritability enrichments of corresponding regions between ohnologs and non-ohnologs

We took advantage of the stratified LD score regression (LDSC) (***Finucane et al., 2015***) to partition heritability 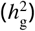 of complex traits among corresponding regions of ohnologs and non-ohnologs. LDSC calculates complex trait’s 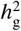 enrichment of genomic regions, (proportion of 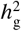)/(proportion of SNPs), using a GWAS result of the complex trait and LD scores of an LDSC model calculated from a reference population. We made an LDSC model from the baseline-LD model that was robust to a wide variety of LD patterns (***Finucane et al., 2015***; ***Gazal et al., 2017***). Our LDSC model, named baselineLD-ohnolog, included annotations of coding regions, introns, UTR and gene flanking regions (0-5 kb, 5-20 kb, and 20-100 kb) (***Yao et al***., 2020) of ohnologs and non-ohnologs (***Singh and*** Isambert, 2020) (refer to Supplementary Table 5 for gene lists and refer to Supplementary Table 6 for annotations).

We applied LDSC to 38 independent complex traits with the baselineLD-ohnolog model (GWAS average *N* = 237, 779; refer to Supplementary Table 1 for a list of traits and refer to Supplementary Materials and Methods for trait selection details). As expected, in coding regions, the 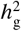 enrichment of ohnologs (random-effects meta-analysis *mean* = 7.7, *se* = 0.50) was significantly higher than of non-ohnologs (*mean* = 4.1, *se* = 0.39; Benjamini-Hochberg = 3.0 × 10^−4^, Wilcoxon signed-rank test; Figure 1A). For both UTR and introns, there was no significant difference between complex traits’ 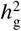 enrichments of ohnologs and non-ohnologs (Figure 1B and 1C). For gene flanking regions, complex traits’ 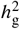 enrichments of ohnologs’ 0-5 kb flanking regions (*mean* = 2.6, *se* = 0.25) were significantly higher than of non-ohnologs’ 0-5 kb flanking regions (*mean* = 1.1, *se* = 0.15; = 6.9 × 10^−5^; Figure 1D); the significance of the difference was reduced in 5-20 kb flanking regions (= 0.040; Figure 1E); the difference was no longer significant in 20-100 kb flanking regions (= 0.63; Figure 1F).

**Figure 1.**
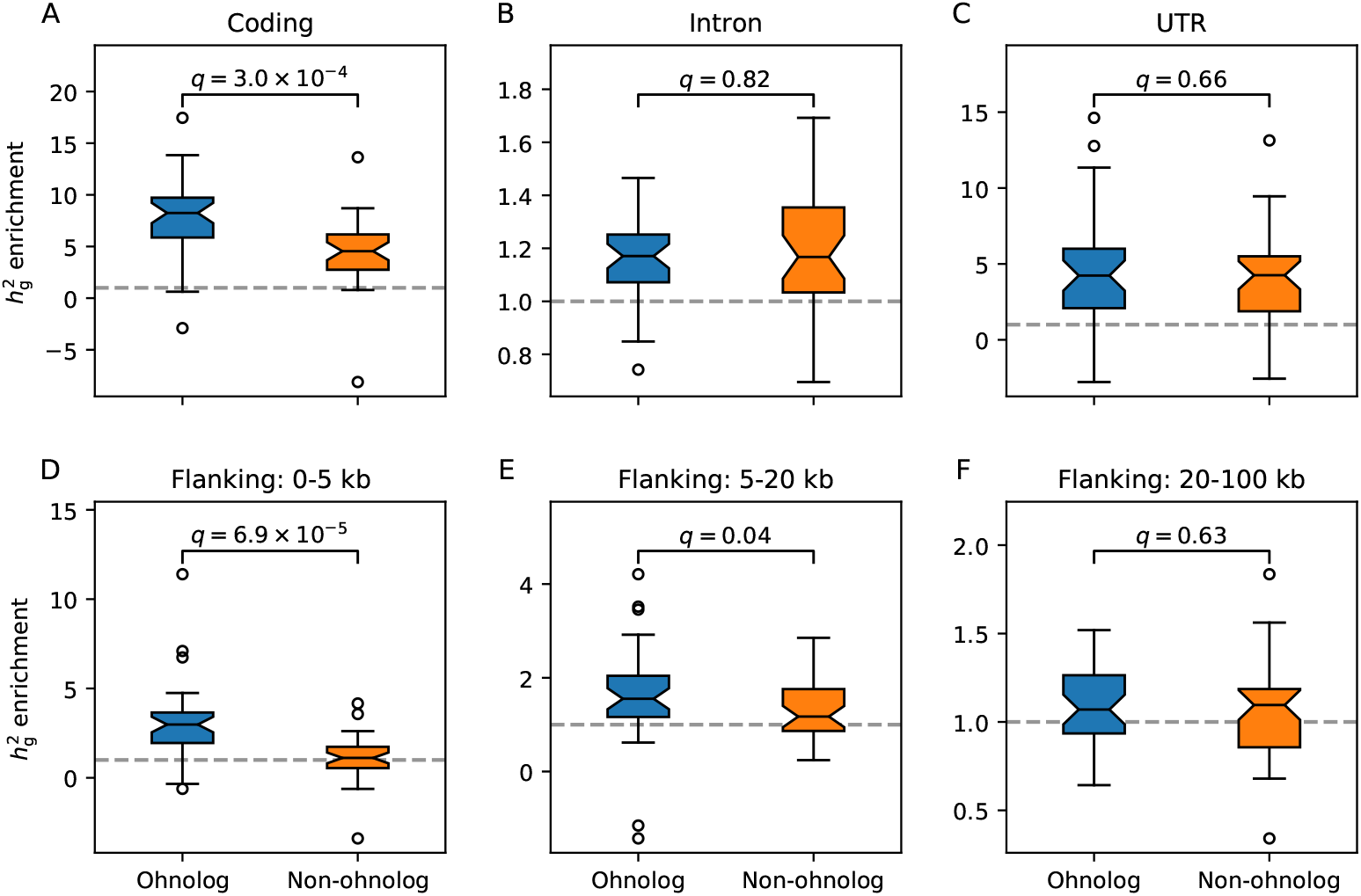
Comparison of 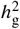 enrichments between ohnologs and non-ohnologs in coding regions (A), intragenic regions (B), untranslated regions (C), 0-5 kb flanking regions (D), 5-20 kb flanking regions (E) and 20-100 kb flanking regions (F). Gray horizontal lines (y=1) indicate no enrichment. Q-values are of Wilcoxon signed-rank tests (*N* = 38) following Benjamini-Hochberg procedure accounting for the six tests. Refer to Supplementary Figure 1-6 for the results of each trait. Refer to Supplementary Table 7 for numeric results. Refer to Supplementary Table 2 for results of random-effects meta-analyses.

### Comparison of complex traits’ expression-mediated heritability enrichments between ohnologs and non-ohnologs

Regulation of gene expression is an essential aspect of the functions of regulatory sequences. In the previous section, we observed that in 5 kb flanking regions of protein-coding genes, the complex traits’ heritability enrichments of ohnologs were significantly higher than of non-ohnologs. Next, we took advantage of the mediated expression score regression (MESC) (***Yao et al***., ***2020***) to quantify the expression-mediated heritability 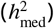 between ohnologs and non-ohnologs, and further test if complex traits’ 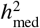 enrichments, 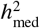 /(proportion of expressed genes), of ohnologs were significantly higher than of non-ohnologs.

For each tissue group, for traits with significantly positive 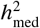 (Benjamini-Hochberg *<* 0.05, one-tailed Z-test, Supplementary Figure 7), we partitioned the 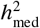 and compared 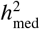 enrichments between ohnologs and non-ohnologs. We observed that the 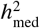 enrichments of ohnologs (random-effects meta-analysis *mean* = 2.1, *se* = 0.12 for the group of all tissues) were significantly higher than of non-ohnologs (*mean* = 0.82, *se* = 0.063; = 8.5 × 10^−5^ for the group of all tissues, Wilcoxon signed-rank test; Figure 2A). As 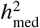 is the product of gene expressio*cis*-heritability 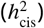 and causal gene expression effect size (***Yao et al***., ***2020***), we further compared 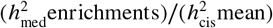, the causal gene expression effect, between ohnologs and non-ohnologs. As 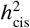 mean of ohnologs was lower than non-ohnologs, when comparing 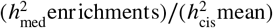 between ohnologs and non-ohnologs, the degree of significance was exacerbated in all tissue groups. The q-values of 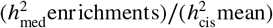 comparisons (Figure 2B) were smaller than the corresponding q-values of 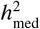 enrichments comparisons (Figure 2A).

**Figure 2.**
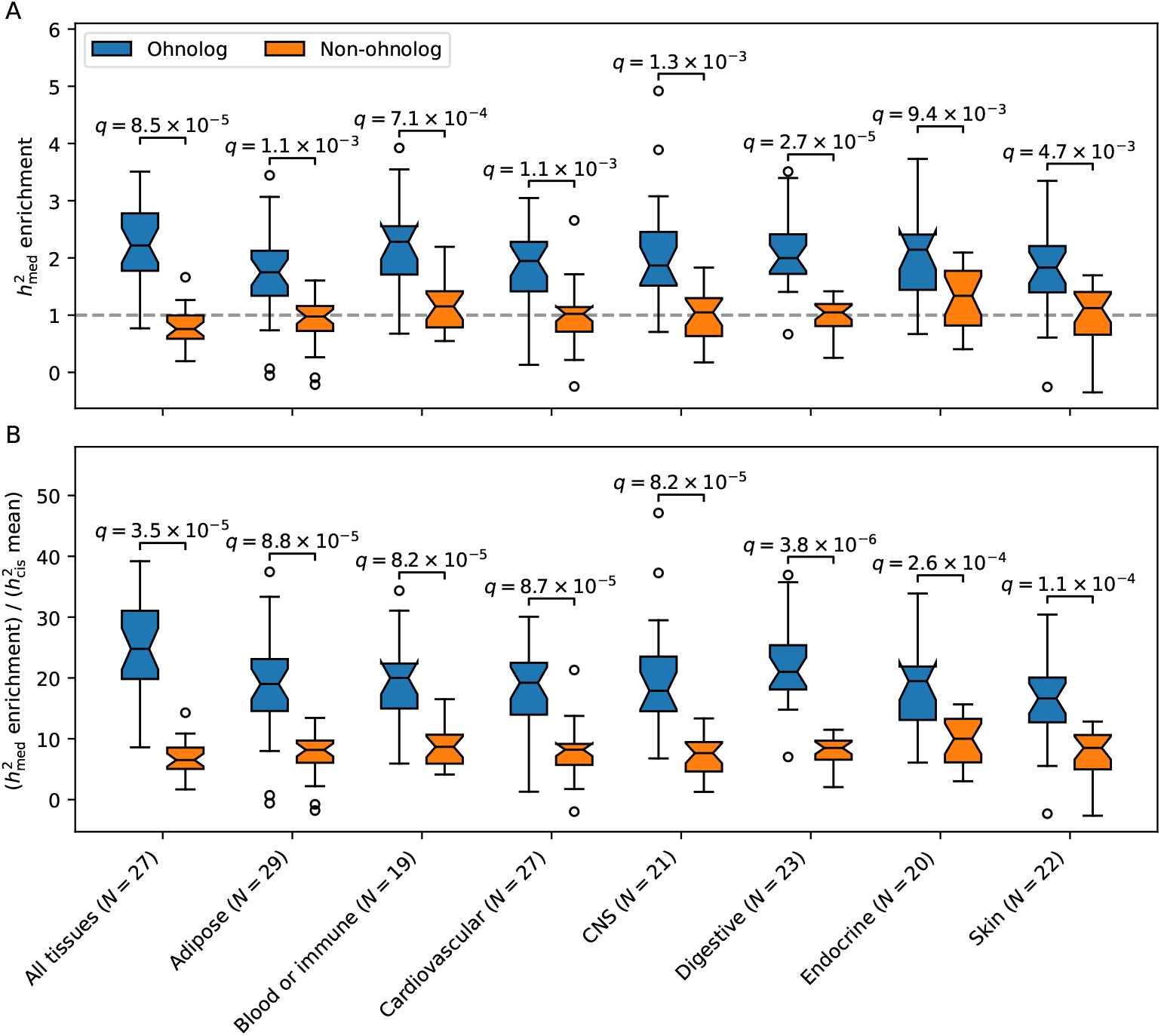
Comparison of 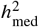 enrichments (A) and causal gene expression effect sizes, 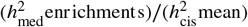 (B), between ohnologs and non-ohnologs. For each tissue group, only traits with significantly positive 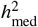 are included. N denotes sample sizes. Q-values are of Wilcoxon signed-rank tests following Benjamini-Hochberg procedure accounting for the eight tests. Refer to Supplementary Figure 8-15 for the results of each trait. Refer to Supplementary Table 8 for numeric results. Refer to Supplementary Table 3 for results of random-effects meta-analyses.

## Discussion

We showed that (i) complex traits’ 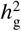 enrichments of regulatory sequences of ohnologs were significantly higher than of non-ohnologs (= 6.9 × 10^−5^ for 0-5 kb gene flanking regions; Figure 1D); (ii) complex traits’ 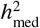 enrichments of ohnologs were significantly higher than of non-ohnologs (= 8.5 × 10^−5^ for the group of all tissues; Figure 2A). To make a further conclusion, we took advantage of Eyre-Walker’s model (***Eyre-Walker,2010***; ***Keightley and Hill, 1990***), which hypothesized that mutations’ absolute values of effect sizes and selection coefficients were positively correlated, either because the trait was a component of fitness or because the mutations had pleiotropic effects on fitness. Recent empirical studies have supported the hypothesis, as heritability of complex traits is more enriched in genes under stronger natural selection regardless of whether a trait affects fitness (***Gazal et al., 2017***; ***Yao et al., 2020***; ***Gazal et al., 2018***). Under Eyre-Walker’s model, we deduce that the natural selection on regulatory sequences of ohnologs is stronger than that of non-ohnologs.

Compared with previous studies, we showed that complex traits’ 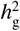 enrichments of coding sequences of ohnologs were significantly higher than of non-ohnologs (*q* = 3.0 × 10^−4^; Figure 1A). Under Eyre-Walker’s model, we deduce that the natural selection on coding sequences of ohnologs is stronger than that of non-ohnologs, which is consistent with the ohnolog retention model of natural selection on coding sequences (***Singh et al., 2012***, ***2014***; ***Roux et al., 2017***). We also showed that complex traits’ causal gene expression effect sizes of ohnologs were significantly larger than of non-ohnologs (*q* = 3.5 × 10^−5^ for the group of all tissues; Figure 2B). Because both regulatory sequences and gene copy number affect gene expression level (***Stranger et al., 2007***), our result illustrates a scenario, natural selection on gene expression level, for the ohnolog retention model of natural selection on gene dosages (***Makino and McLysaght, 2010***; ***McLysaght et al., 2014***; ***Rice*** and McLysaght, 2017).

In conclusion, previous studies showed that ohnologs were vulnerable to coding sequence mutations (***Singh et al., 2012***, ***2014***; ***Roux et al., 2017***) and copy numbers mutations (***Makino and*** McLysaght, 2010; ***McLysaght et al., 2014***; ***Rice and McLysaght, 2017***). We showed that ohnologs were also vulnerable to regulatory sequence mutations. We believe that both amino acid sequences and expression levels of ohnologs are under substantial natural selection. The natural selection on regulatory sequences ensures that ohnologs are stably expressed, thus maintain a functional state.

## Methods and Materials

We curated GWAS results of 38 independent complex traits and diseases (***Abbott et al***., 2018; ***De***montis et al., 2019; ***Watson et al***., 2019; ***Hujoel et al., 2020***; ***de Lange et al., 2017***; ***Nalls et al., 2019***; ***Okada et al., 2014***; ***Pardiñas et al., 2018***; ***Mahajan et al., 2018***; ***Yu et al., 2019; Jin et al., 2016***) with the standard of independence described by ***Finucane et al. (***2015***)*** (Supplementary Table 1 and 4). We used the Ensembl GRCh37 annotated autosomal protein-coding genes as the universal gene set. We used the human ohnolog list generated by the intermediate criterion of ***Singh and*** Isambert (***2020***) (Supplementary Table 5). We made the baselineLD-ohnolog model by adding annotations of coding regions, introns, UTR and gene flanking regions (0-5 kb, 5-20 kb, and 20-100 kb) of ohnologs and non-ohnologs into the baseline-LD model v2.0 (***Gazal et al***., 2017) and excluding linearly dependent annotations (Supplementary Table 6). We applied stratified LDSC to the 38 independent traits using the baselineLD-ohnolog model as described previously (***Gazal et al***., 2017). The expression scores of ohnologs and non-ohnologs were calculated from the eQTL effect sizes of GTEx v8 tissues (***Yao et al., 2020***; ***GTEx, 2017***) with tissue groups defined by ***Yao et al.*** (***2020***). We applied stratified MESC on traits with significantly positive 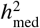 (Supplementary Figure 7-15) for each tissue group to calculate 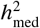 enrichments of ohnologs and non-ohnologs. We added two annotations of all SNPs within 100 kb of ohnologs and non-ohnologs in addition to the baseline-LD model to ensure the elimination of false positive 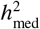 enrichment (***Yao et al***., 2020) when performing MESC. For a comparison of 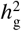 or 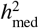 enrichments between ohnologs and non-ohnologs, as each trait had values of ohnologs and non-ohnologs, we performed a Wilcoxon signed-rank test. We used the Benjamini-Hochberg method to calculate q-values. We used the Paule and Mandel method to perform the random-effects meta-analysis of 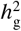 or 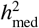. All statistical analyses were performed using the Python package of Statsmodels v0.12 (***Seabold and Perktold, 2010***). Refer to Supplementary Materials and Methods for more details.

## Acknowledgments

We acknowledge Chao Xue for help in collating GWAS data. We acknowledge Dr. Bolan Yu for help during the COVID-19 pandemic. This work was funded by National Key R&D Program of China (2018YFC0910500) and National Natural Science Foundation of China (31771401).

