## Supplementary Materials and Methods, Figures and Tables 1-3 for "Natural selections on both amino acid sequences and expression levels are determinants of ohnolog retention"

*Complex traits and diseases*

The GWAS results of UKBB traits produced by Dr. Benjamin Neale's lab (Abbott et al. 2018) were downloaded. We chose GWAS results (n=136) that met all of the following conditions for subsequent analyses: (1) was "primary" GWAS, (2) the confidence was "high", (3) Z-score of non-zero heritability was greater than seven and (3) the observed heritability was greater than 0.1. Then we removed traits to get a set of independent traits with the standard described by Finucane et al. (2015):

> "For our meta-analysis over traits, we identified pairs of traits with substantial sample overlap and trait correlation by using the intercept of cross-trait LD score regression (Bulik-Sullivan, Finucane, et al. 2015). Specifically, for each pair of traits, we computed the genetic covariance intercept on the N1N2 scale, which for quantitative traits estimates phenotypic correlation times sample overlap, and for case-control estimates a related quantity that is high, for example, if two unrelated traits share controls. This intercept is downwardly biased in the presence of GC correction, so we divided by the square root of the product of the heritability estimates of the two traits to correct for this bias. We identified pairs of traits for which this quantity was at least 15% of the sample size of either of the traits, and we excluded one of each such pair."

We used an iterative procedure to get as many independent traits as possible: (1) for each trait, calculate the number (n) of other traits correlated to the trait; (2) from traits with the largest n, remove the trait with smallest Z-value of non-zero heritability; (3) repeat (1) and (2) until all traits were independent. We also curated GWAS results of complex diseases with a Z-score of non-zero heritability greater than six. After merging the independent disease traits with the independent UKBB traits and removing correlated traits, 38 independent traits finally remained (Okada et al. 2014; Jin et al. 2016; de Lange et al. 2017; Abbott et al. 2018; Mahajan et al. 2018; Pardiñas et al. 2018; Demontis et al. 2019; Nalls et al. 2019; Watson et al. 2019; Yu et al. 2019; Hujoel et al. 2020) (Supplementary Table 1 and 4).

*Gene lists of ohnologs and non-ohnologs*

We downloaded the human ohnolog list generated with Singh's intermediate criterion from the OHNOLOGS v2 website (Singh and Isambert 2020). We took an intersection between the set of Singh’s ohnolog list (n=5,954) and the set of Ensembl GRCh37 autosomal protein-coding genes (n=18,336) to get a set of autosomal protein-coding ohnologs (n=5,528, we referred to it just as ohnologs for short). The non-ohnolog gene set (n=12,808) was the subtraction of the ohnolog gene set from the set of Ensembl GRCh37 autosomal protein-coding genes (Supplementary Table 5).

*LDSC*

The original baseline model is robust to a wide variety of enrichment patterns because including many categories gives it the flexibility to adapt to the unknown causal model (Finucane et al. 2015). The baseline-LD model, including more categories, is more robust to different LD patterns (Gazal et al. 2017). We downloaded the baseline-LD model v2.0, and the quality controlled PLINK files of 1000 Genomes Project European population (Gazal et al. 2017). SNP annotations of ohnologs and non-ohnologs were generated using the make_annot.py script of the LDSC package (Finucane et al. 2015) with 0 kb, 5 kb, 20 kb and 100 kb extensions respectively. The annotation of ohnolog coding regions was the intersection of the ohnolog (0 kb extension) annotation and the baseline-LD annotation of UCSC coding. The annotation of ohnolog intron was the intersection of the ohnolog (0 kb extension) annotation and the baseline-LD annotation of UCSC intron. The annotation of ohnolog UTR was the intersection of the ohnolog (0 kb extension) annotation and the union of baseline-LD annotations of UCSC 3' UTR and UCSC 5'UTR. The annotation of ohnolog 0-5 kb flanking regions was subtracting the ohnolog (0 kb extension) annotation from the ohnolog (5 kb extension) annotation. The annotation of ohnolog 5-20 kb flanking regions was subtracting the ohnolog (5 kb extension) annotation from the ohnolog (20 kb extension) annotation. The annotation of ohnolog 20-100 kb flanking regions was subtracting the ohnolog (20 kb extension) annotation from the ohnolog (100 kb extension) annotation. Corresponding procedures produced the corresponding annotations of non-ohnologs. The non-gene annotation was subtracting the ohnolog (100 kb extension) and the non-ohnolog (100 kb extension) from the baseline-LD base annotation. We excluded the baseline-LD annotations of UCSC coding, UCSC intron, UCSC 3'UTR, UCSC 5'UTR and base because they were linearly dependent on the newly added annotations. We named the model baselineLD-ohnolog, which contained 84 annotations (Supplementary Table 6). Then, LD scores were calculated using the quality controlled PLINK files of 1000 Genomes Project European population (Gazal et al. 2017). Finally, we applied LDSC to the 38 independent traits with the baselineLD-ohnolog. The LDSC was performed on HapMap 3 SNPs without MHC region, and heritability was calculated using SNPs with MAF≥5%.

*MESC*

The expression scores and eQTL effect sizes of GTEx v8 individual tissues were downloaded from MESC website (Yao et al. 2020). The gene lists of ohnologs and non-ohnologs described above were used to generate the gene set expression scores. The expression scores of all genes and gene sets were calculated by meta_analyze_weights.py and gene_set_analysis.py scripts of the MESC package respectively with tissue groups defined by Yao et al. (2020) and with the quality controlled PLINK files of 1000 Genomes Project European population (Gazal et al. 2017). We estimated the overall $h^2_{\mathrm{med}}$ of all tissue groups by MESC. For each tissue group, for traits with positive $h^2_{\mathrm{med}}$ estimates, we tested if $h^2_{\mathrm{med}}$ was significantly larger than zero with a one-tailed Z-test. For each tissue group, for traits with q<0.05 of positive $h^2_{\mathrm{med}}$, we further estimated $h^2_{\mathrm{med}}$ enrichments of ohnologs and non-ohnologs by MESC. When estimating $h^2_{\mathrm{med}}$ enrichments of ohnologs and non-ohnologs, in addition to the baseline-LD model, we added two annotations of all SNPs within 100 kb of ohnologs or non-ohnologs, which could eliminate false positive $h^2_{\mathrm{med}}$ enrichment estimates that arose due to localization of GWAS signal around genes that was not mediated by gene expression (Yao et al. 2020).

*URLs*

Human ohnolog list (Singh and Isambert 2020), http://ohnologs.curie.fr/;

GWAS results of UK Biobank traits (Abbott et al. 2018), http://www.nealelab.is/uk-biobank;

LDSC software (Bulik-Sullivan, Loh, et al. 2015), http://www.github.com/bulik/ldsc/;

Quality controlled PLINK files of 1000 Genomes Project European Population and baseline-LD model v2.0 (Gazal et al. 2017), http://alkesgroup.broadinstitute.org/LDSCORE/;

MDSC software and expression scores (Yao et al. 2020), http://github.com/douglasyao/mesc/.

**Supplementary Tables** (Refer to the Excel file for Supplementary Table 4-8)

Supplementary Table 1. Traits subjected to LDSC and MESC

Supplementary Table 2. Random-effect meta-analyses of proportion and enrichment of $h_g^2$

Supplementary Table 3. Random-effect meta-analyses of proportion and enrichment of $h_{med}^2$

Supplementary Table 4. Genetic correlation matrix of traits subjected to LDSC and MESC

Supplementary Table 5. Gene lists of ohnologs and non-ohnologs

Supplementary Table 6. Annotations of the baselineLD-ohnolog model

Supplementary Table 7. Numeric results of stratified LDSC

Supplementary Table 8. Numeric results of stratified MESC

**Supplementary Figures**

Supplementary Figure 1-6. Comparison of $h_g^2$ enrichments between ohnologs and non-ohnologs in coding regions (Supplementary Figure 1), intragenic regions Supplementary Figure 2), untranslated regions (Supplementary Figure 3), 0-5 kb flanking regions (Supplementary Figure 4), 5-20 kb flanking regions (Supplementary Figure 5) and 20-100 kb flanking regions (Supplementary Figure 6) for each trait. Gray vertical lines (x=1) indicate no enrichment.

Supplementary Figure 7. Comparison of Z-values of positive $h_{med}^2$ among tissue groups for each trait. Asterisks indicate significantly positive $h_{med}^2$ (q<0.05, one-tailed Z-test).

Supplementary Figure 8-15. Each figure shows $h^2_{\text{med}}/h^2_{\text{g}}$ (left) and $h^2_{\text{med}}$ enrichments of ohnologs and non-ohnologs (right) of one tissue group. Q-values are of one-tailed Z-tests of positive $h^2_{\text{med}}$.

Supplementary Table 1. Traits subjected to LDSC and MESC

| Phenotype | Observed $h^2_g$ | $h^2_g$ $Z$-value | Sample size | Reference | Description | Abbreviation |
|---|---|---|---|---|---|---|
| Pulse rate | 0.1571 | 15.226 | 340,162 | Abbott et al. 2018 | Pulse rate, automated reading | Pulse |
| Chronotype | 0.1187 | 21.971 | 322,488 | Abbott et al. 2018 | Morning/evening person (chronotype) | Chronotype |
| Alcohol preference | 0.1012 | 16.339 | 184,716 | Abbott et al. 2018 | Alcohol usually taken with meals | Alcohol |
| Fluid intelligence | 0.2225 | 19.272 | 117,131 | Abbott et al. 2018 | Fluid intelligence score | Fluid |
| Neuroticism | 0.1119 | 13.560 | 293,006 | Abbott et al. 2018 | Neuroticism score | Neurotic |
| Impedance of arm | 0.2482 | 30.855 | 354,792 | Abbott et al. 2018 | Impedance of arm (right) | Impedance |
| Age of facial hair | 0.1330 | 13.816 | 161,470 | Abbott et al. 2018 | Relative age of first facial hair | Beard |
| Balding pattern 1 | 0.2277 | 10.615 | 165,649 | Abbott et al. 2018 | Hair/balding pattern: Pattern 1 | Balding |
| Age of menarche | 0.2090 | 17.338 | 188,644 | Abbott et al. 2018 | Age when periods started (menarche) | Menarche |
| Child birth weight | 0.1115 | 14.909 | 155,202 | Abbott et al. 2018 | Birth weight of first child | First |
| Age at last birth | 0.1046 | 13.994 | 131,806 | Abbott et al. 2018 | Age at last live birth | Last |
| RBC | 0.2337 | 10.044 | 350,475 | Abbott et al. 2018 | Red blood cell (erythrocyte) count | RBC |
| RDW | 0.2172 | 12.178 | 350,473 | Abbott et al. 2018 | Red blood cell (erythrocyte) distribution width | RDW |
| Eosinophill count | 0.1840 | 13.869 | 349,856 | Abbott et al. 2018 | Eosinophill count | EOS |
| Lymphocyte percentage | 0.1632 | 12.888 | 349,861 | Abbott et al. 2018 | Lymphocyte percentage | Lymphocyte |
| IRF | 0.1641 | 7.656 | 344,728 | Abbott et al. 2018 | Immature reticulocyte fraction | IRF |
| FVC | 0.2100 | 25.756 | 329,404 | Abbott et al. 2018 | Forced vital capacity (FVC) | FVC |
| Creatinine | 0.2115 | 16.803 | 344,104 | Abbott et al. 2018 | Creatinine (umol/L) | Creatinine |
| IGF-1 | 0.2532 | 13.711 | 342,439 | Abbott et al. 2018 | IGF-1 (nmol/L) | IGF1 |
| Phosphate | 0.1336 | 7.857 | 314,658 | Abbott et al. 2018 | Phosphate (mmol/L) | Phosphate |
| Total protein | 0.1666 | 12.930 | 314,921 | Abbott et al. 2018 | Total protein (g/L) | TP |
| Bone density | 0.3175 | 10.195 | 206,496 | Abbott et al. 2018 | Heel bone mineral density (BMD) | BMD |
| Age of menopause | 0.1148 | 8.975 | 111,593 | Abbott et al. 2018 | Age at menopause (last menstrual period) | Menopause |
| Spherical power | 0.3264 | 16.241 | 77,983 | Abbott et al. 2018 | Spherical power (right) | Spherical |
| 6mm strong meridian | 0.4230 | 11.435 | 66,256 | Abbott et al. 2018 | 6mm strong meridian (right) | Meridian |
| IOP | 0.1692 | 11.524 | 76,510 | Abbott et al. 2018 | Intra-ocular pressure, corneal-compensated (left) | IOP |
| CRF | 0.3113 | 11.674 | 76,510 | Abbott et al. 2018 | Corneal resistance factor (left) | CRF |
| ADHD | 0.1674 | 13.392 | 55,374 | Demontis et al. 2019 | Attention deficit hyperactivity disorder | ADHD |
| Anorexia nervosa | 0.1332 | 12.566 | 72,517 | Watson et al. 2019 | Anorexia nervosa | Anorexia |
| Breast cancer | 0.0173 | 7.522 | 427,359 | Hujoel et al. 2020 | Breast cancer | Breast |
| Crohn's disease | 0.3030 | 8.464 | 40,266 | de Lange et al. 2017 | Crohn's disease | Crohn |
| Parkinson's disease | 0.0123 | 8.786 | 482,730 | Nalls et al. 2019 | Parkinson's disease | Parkinson |
| Prostate cancer | 0.0153 | 8.053 | 415,704 | Hujoel et al. 2020 | Prostate cancer | Prostate |
| Rheumatoid arthritis | 0.0964 | 7.248 | 58,284 | Okada et al. 2014 | Rheumatoid arthritis | RA |
| Schizophrenia | 0.3099 | 20.660 | 105,318 | Pardinas et al. 2018 | Schizophrenia | SCZ |
| Type 2 diabetes | 0.1557 | 19.709 | 898,130 | Mahajan et al. 2018 | Type 2 diabetes | T2D |
| Tourette syndrome | 0.2406 | 6.683 | 14,307 | Yu et al. 2019 | Tourette syndrome | Tourette |
| Vitiligo | 0.1197 | 6.470 | 44,266 | Jin et al. 2016 | Vitiligo | Vitiligo |

Supplementary Table 2. Random-effect meta-analyses of proportion and enrichment of $h^2_g$

| | Proportion of common SNPs | Proportion of $h^2_g$ | | $h^2_g$ enrichment | |
|---|---|---|---|---|---|
| | | Mean | SE | Mean | SE |
| Coding | | | | | |
| Ohnolog | 0.49% | 3.80% | 0.24% | 7.7256 | 0.4968 |
| Non-ohnolog | 0.92% | 3.80% | 0.36% | 4.1350 | 0.3891 |
| UTR | | | | | |
| Ohnolog | 0.48% | 2.13% | 0.24% | 4.4672 | 0.5048 |
| Non-ohnolog | 0.83% | 2.89% | 0.35% | 3.4703 | 0.4247 |
| Intron | | | | | |
| Ohnolog | 17.74% | 20.72% | 0.41% | 1.1684 | 0.0229 |
| Non-ohnolog | 18.44% | 21.43% | 0.59% | 1.1619 | 0.0322 |
| Flanking: 0-5 kb | | | | | |
| Ohnolog | 1.60% | 4.17% | 0.39% | 2.6015 | 0.2456 |
| Non-ohnolog | 3.29% | 3.54% | 0.49% | 1.0753 | 0.1496 |
| Flanking: 5-20 kb | | | | | |
| Ohnolog | 4.49% | 6.59% | 0.55% | 1.4679 | 0.1224 |
| Non-ohnolog | 7.06% | 8.16% | 0.60% | 1.1557 | 0.0856 |
| Flanking: 20-100 kb | | | | | |
| Ohnolog | 16.92% | 18.13% | 0.52% | 1.0715 | 0.0307 |
| Non-ohnolog | 18.79% | 19.88% | 0.73% | 1.0580 | 0.0391 |
| Non-gene | 29.31% | 16.85% | 1.13% | 0.5748 | 0.0387 |

Supplementary Table 3. Random-effect meta-analyses of proportion and enrichment of $h^2_{med}$

| | expressed genes | Average $h^2_{cis}$ | Proportion of $h^2_{med}$ | | $h^2_{med}$ enrichment | |
|---|---|---|---|---|---|---|
| | | | Mean | SE | Mean | SE |
| All tissues | | | | | | |
| Non-ohnolog | 44.48% | 11.64% | 36.67% | 2.81% | 0.8245 | 0.0631 |
| Ohnolog | 20.82% | 8.95% | 44.20% | 2.46% | 2.1231 | 0.1183 |
| Adipose | | | | | | |
| Non-ohnolog | 44.69% | 11.95% | 43.55% | 2.78% | 0.9745 | 0.0621 |
| Ohnolog | 21.20% | 9.20% | 34.84% | 2.23% | 1.6436 | 0.1051 |
| Blood or immune | | | | | | |
| Non-ohnolog | 45.60% | 13.29% | 52.55% | 3.43% | 1.1524 | 0.0752 |
| Ohnolog | 20.94% | 11.42% | 38.25% | 3.30% | 1.8264 | 0.1576 |
| Cardiovascular | | | | | | |
| Non-ohnolog | 46.11% | 12.47% | 45.19% | 2.88% | 0.9802 | 0.0624 |
| Ohnolog | 21.81% | 10.14% | 39.33% | 2.55% | 1.8033 | 0.1170 |
| CNS | | | | | | |
| Non-ohnolog | 44.68% | 13.72% | 45.68% | 4.35% | 1.0224 | 0.0975 |
| Ohnolog | 20.48% | 10.44% | 36.72% | 3.05% | 1.7930 | 0.1491 |
| Digestive | | | | | | |
| Non-ohnolog | 45.19% | 12.34% | 45.21% | 2.94% | 1.0004 | 0.0650 |
| Ohnolog | 21.25% | 9.51% | 44.43% | 2.77% | 2.0908 | 0.1303 |
| Endocrine | | | | | | |
| Non-ohnolog | 40.89% | 13.37% | 52.52% | 4.68% | 1.2843 | 0.1145 |
| Ohnolog | 19.14% | 11.01% | 36.67% | 2.91% | 1.9159 | 0.1523 |
| Skin | | | | | | |
| Non-ohnolog | 45.45% | 13.22% | 49.93% | 3.89% | 1.0987 | 0.0856 |
| Ohnolog | 21.87% | 11.00% | 40.14% | 3.59% | 1.8355 | 0.1643 |

Supplementary Figure 1

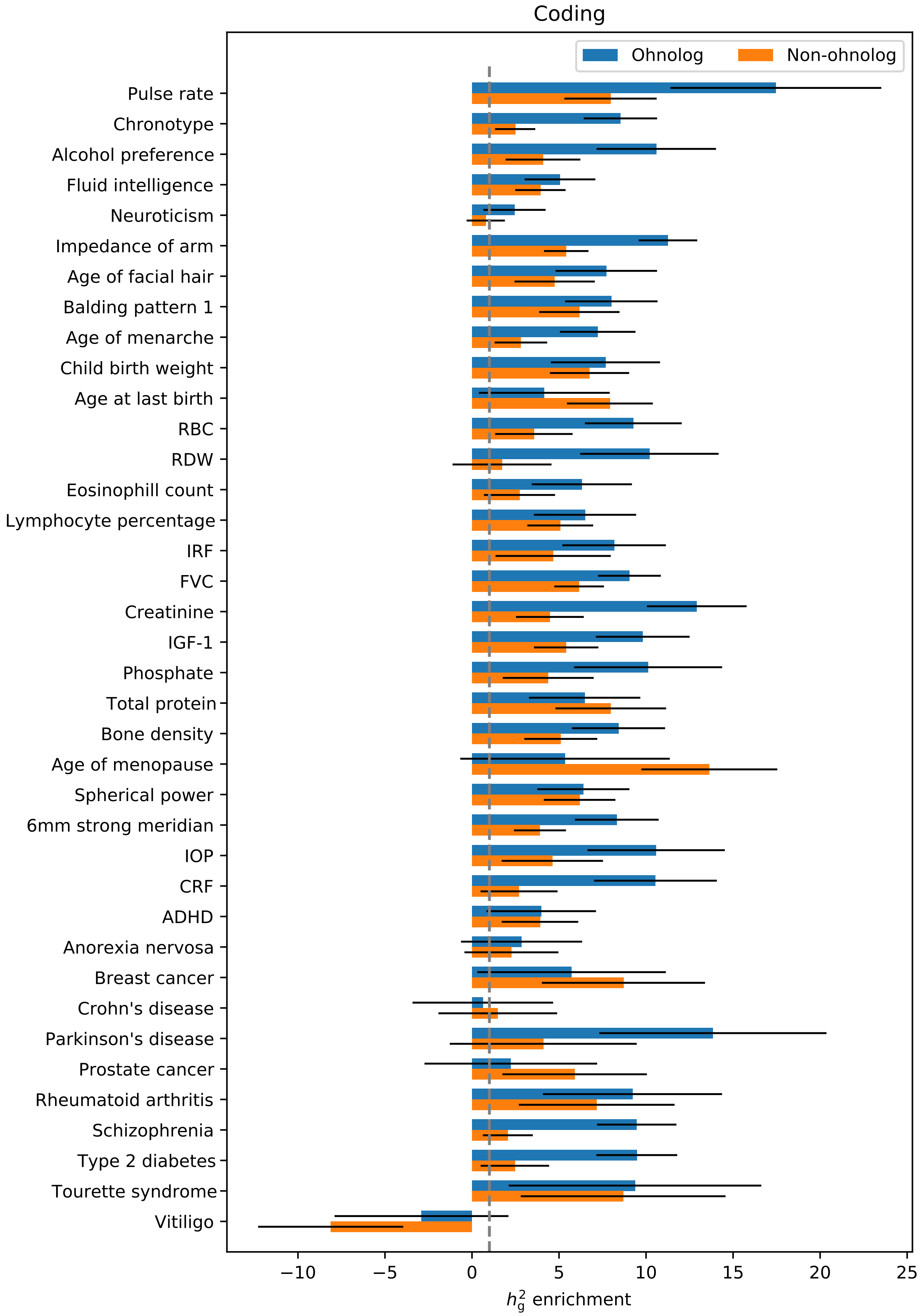

Supplementary Figure 2

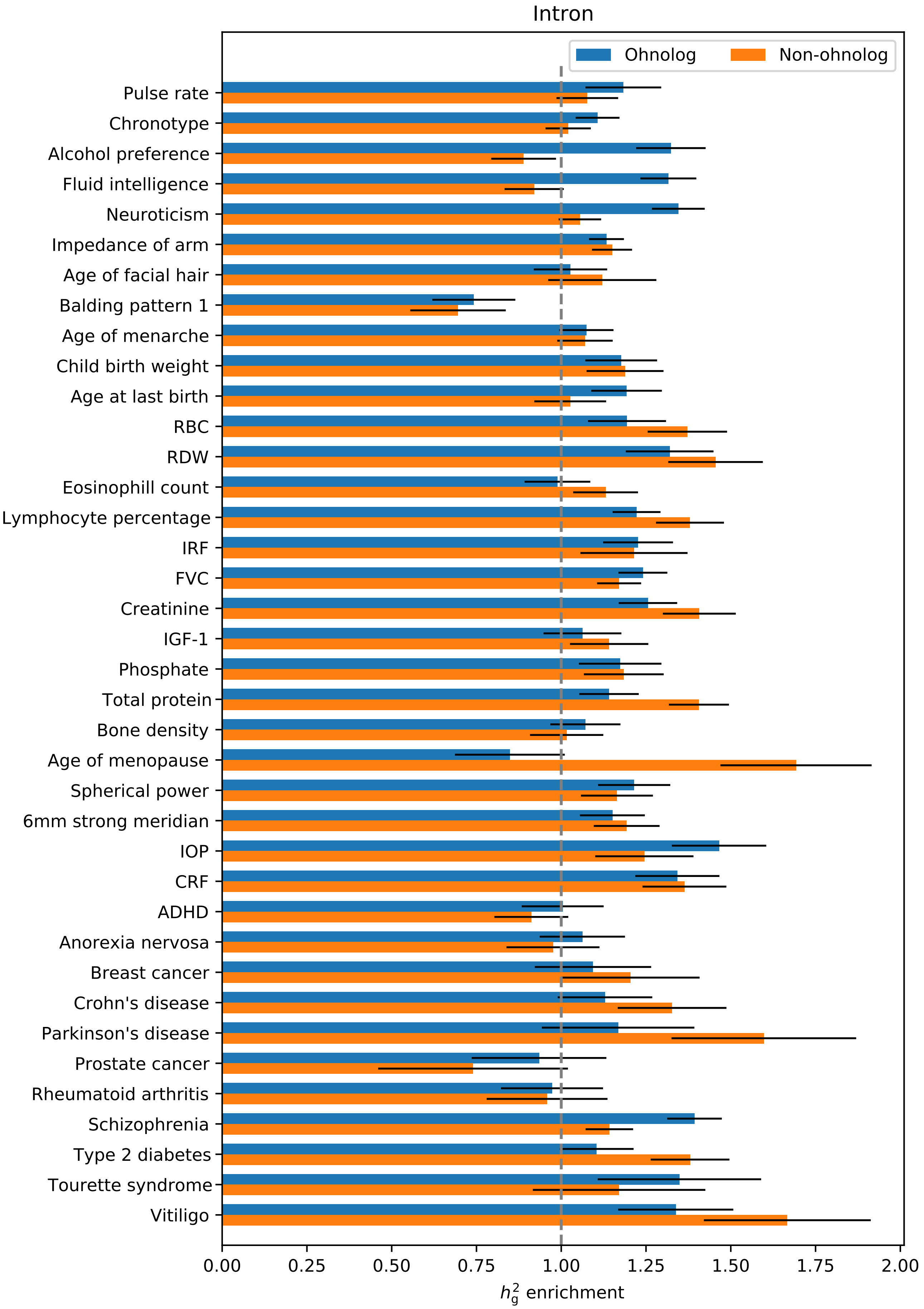

Supplementary Figure 3

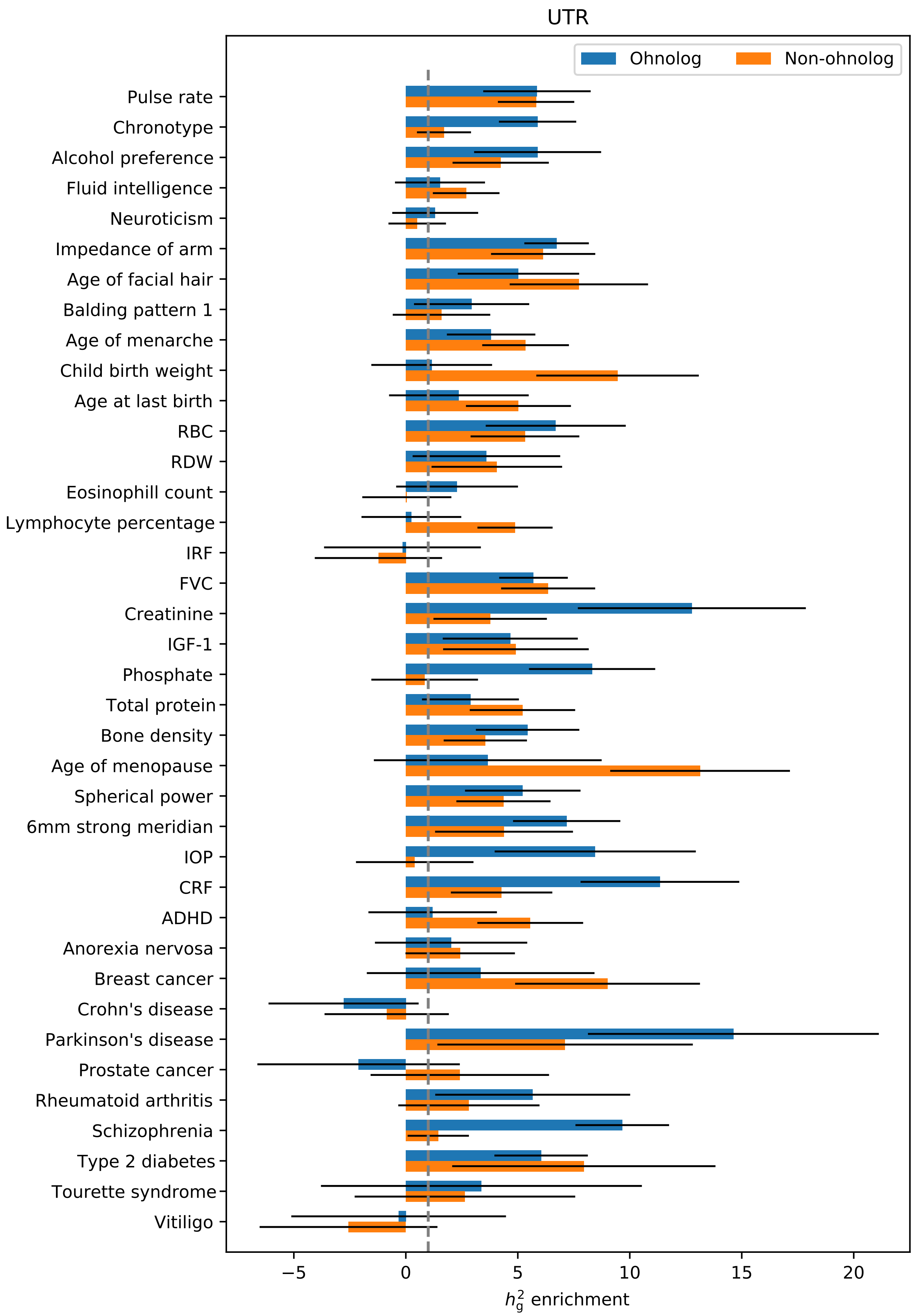

Supplementary Figure 4

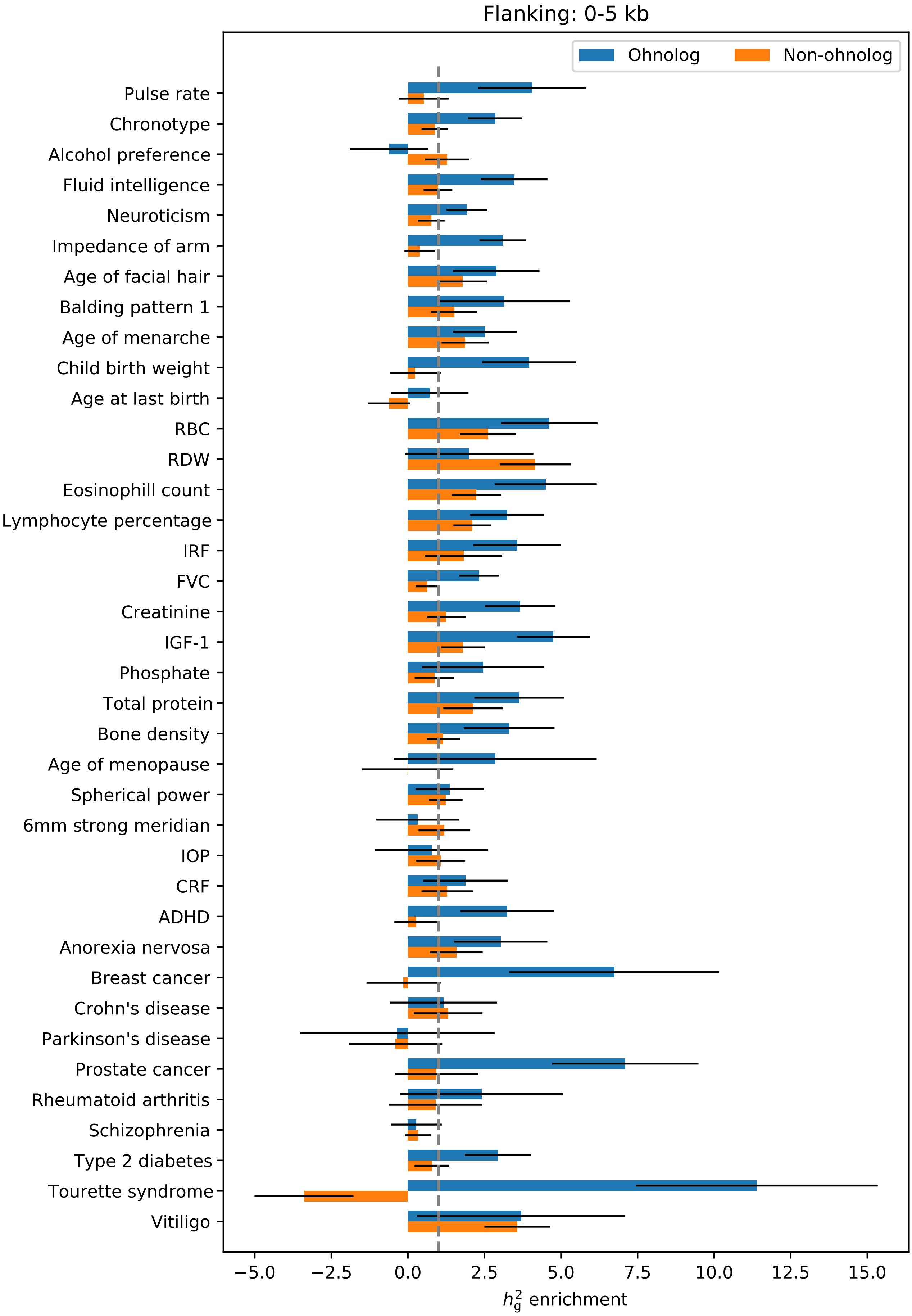

Supplementary Figure 5

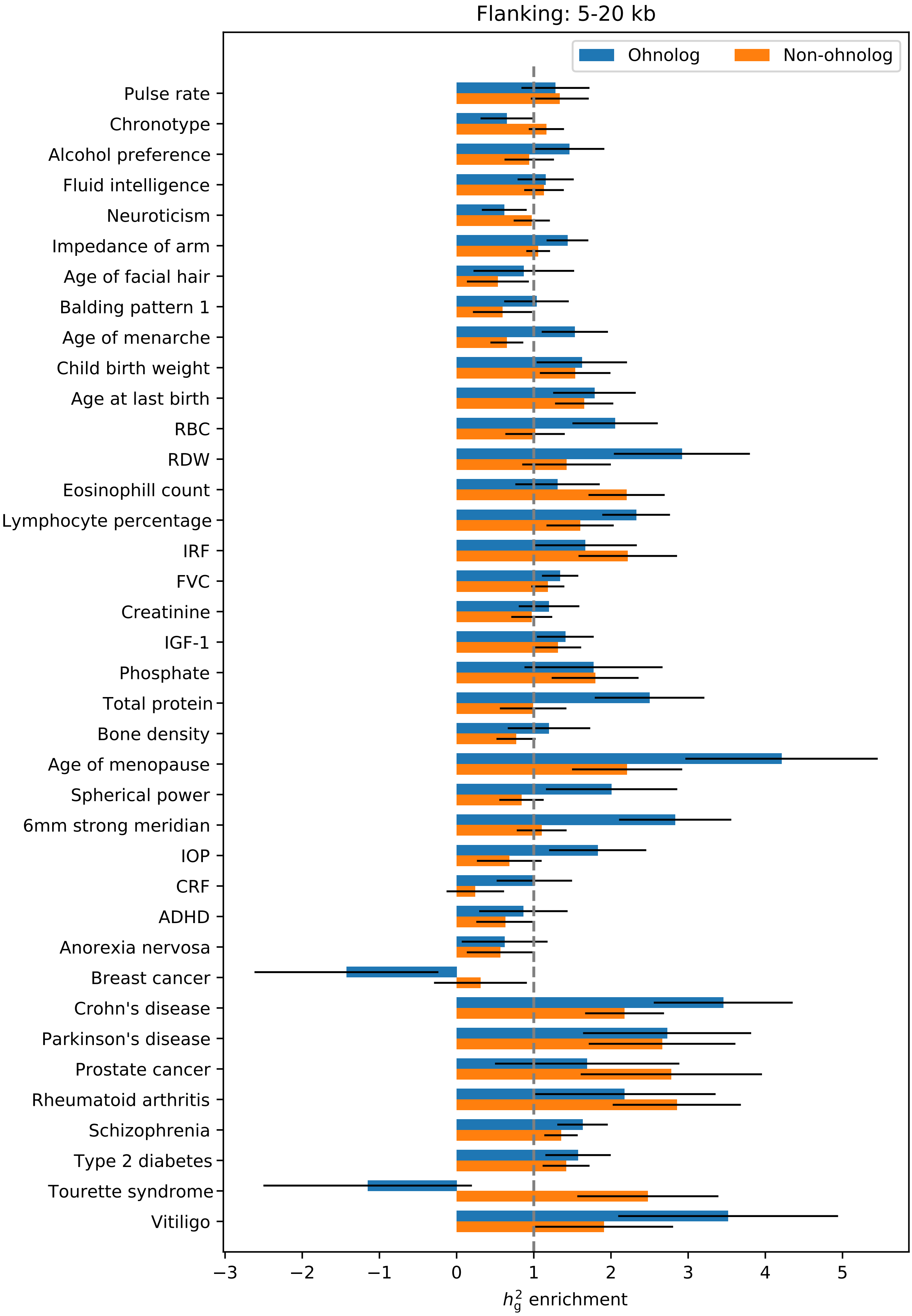

Supplementary Figure 6

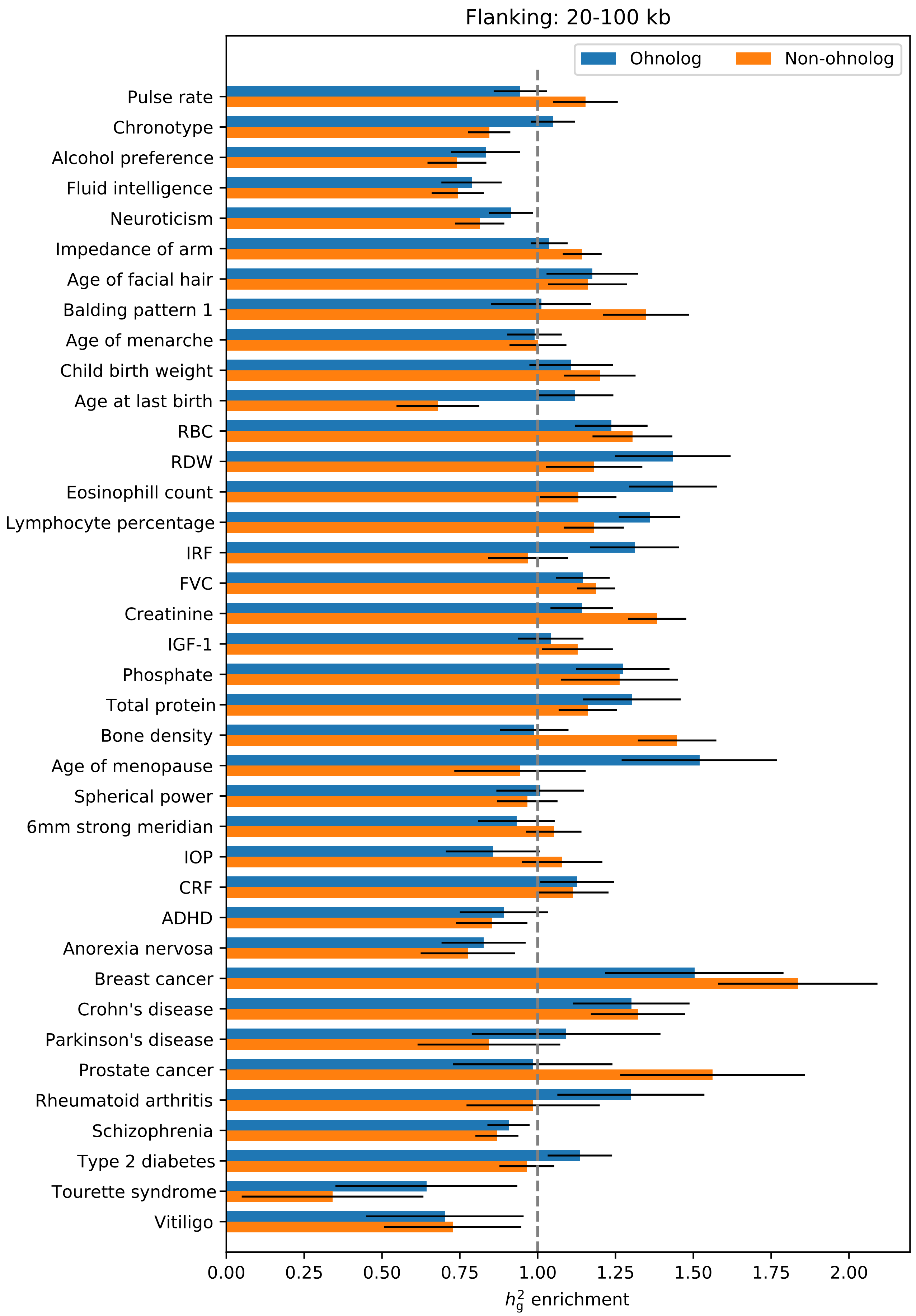

Supplementary Figure 7

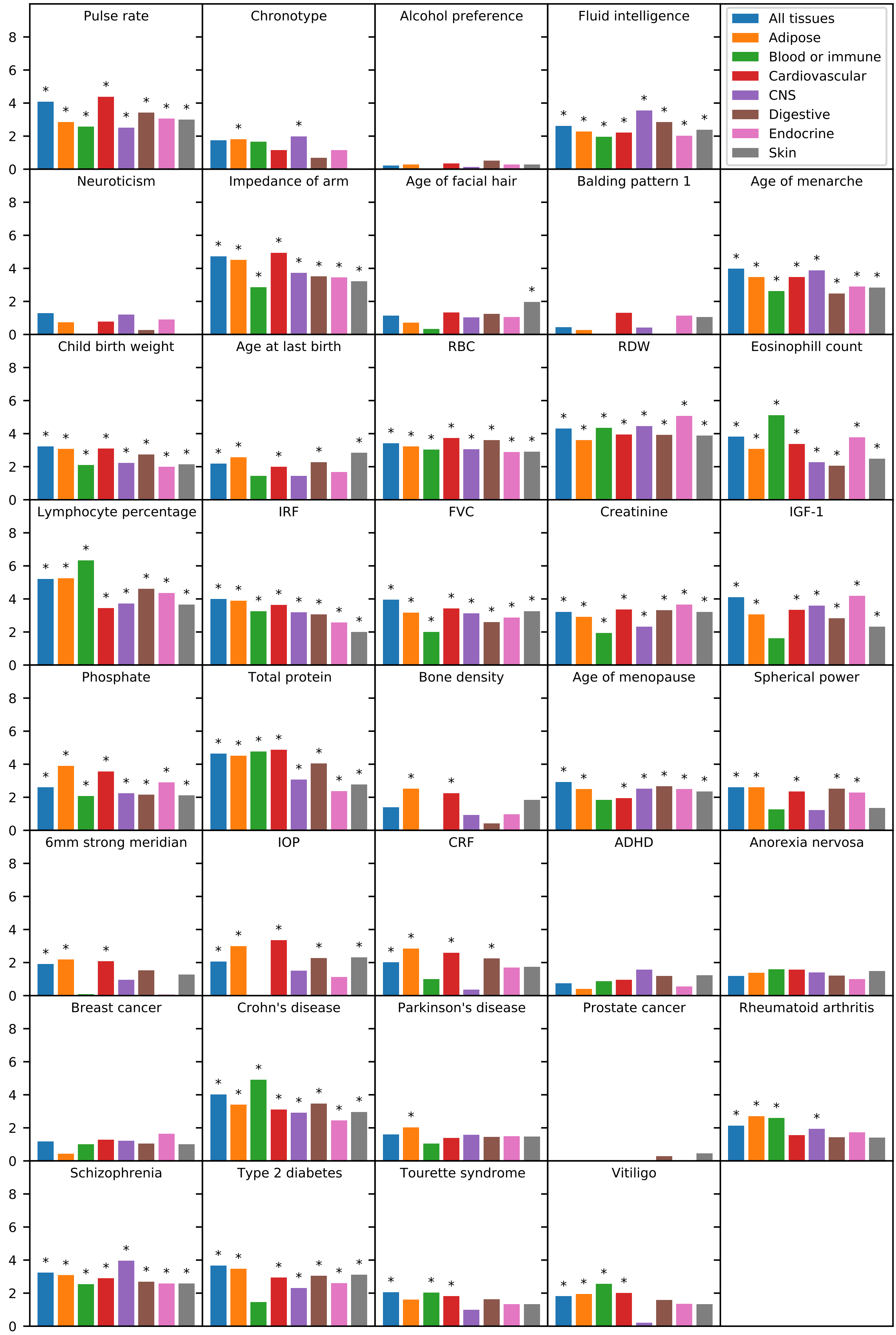

Supplementary Figure 8

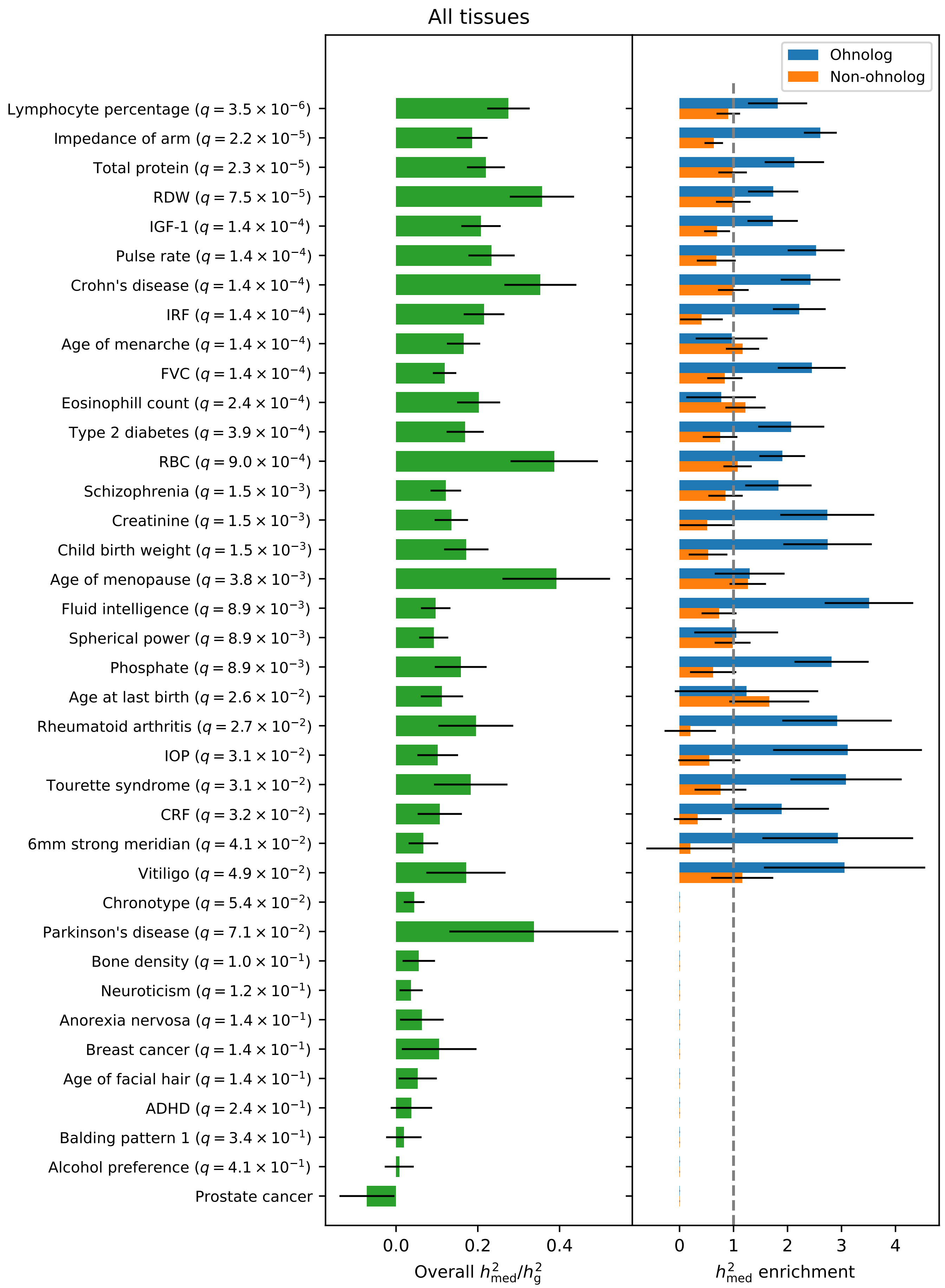

Supplementary Figure 9

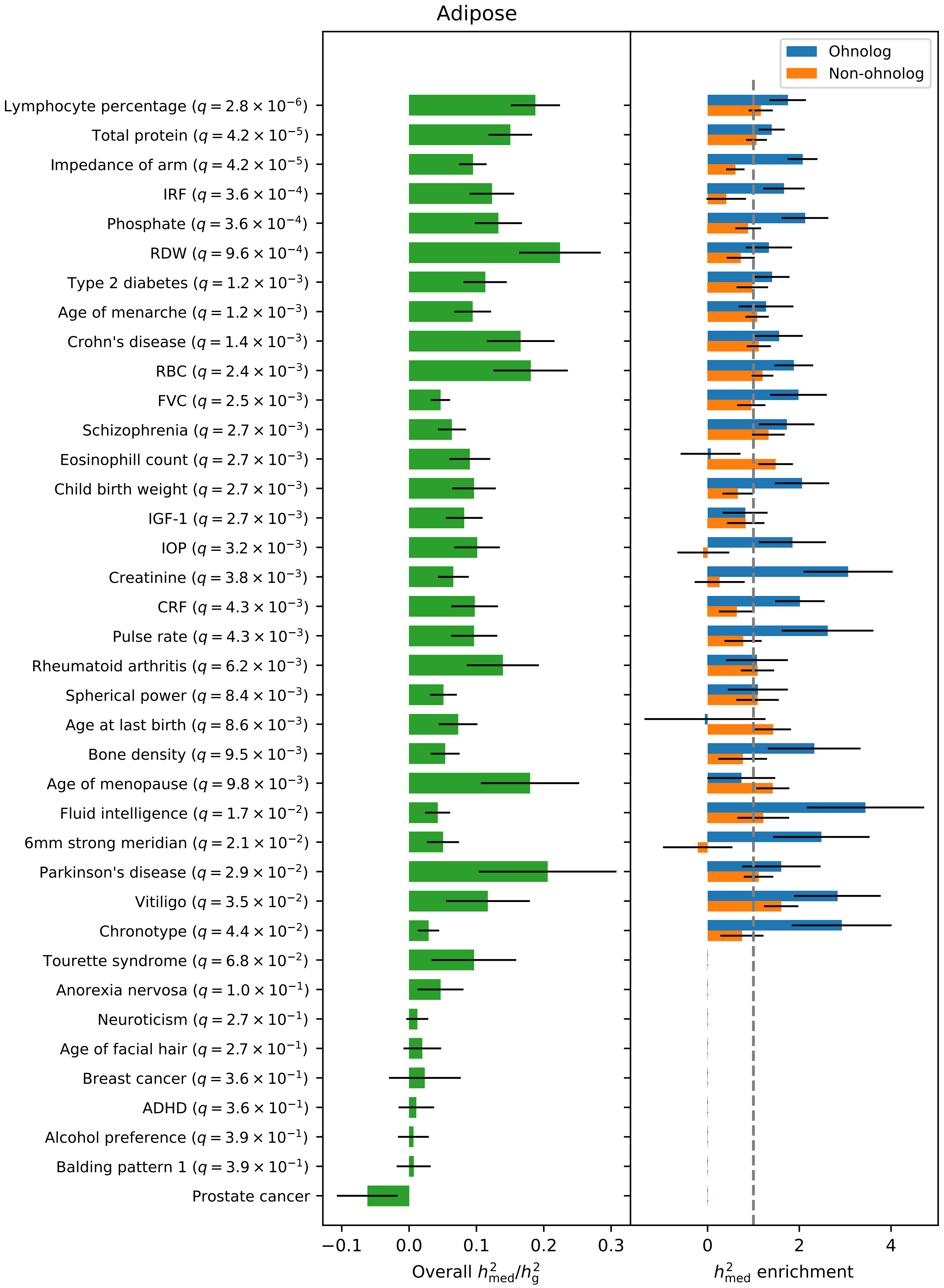

Supplementary Figure 10

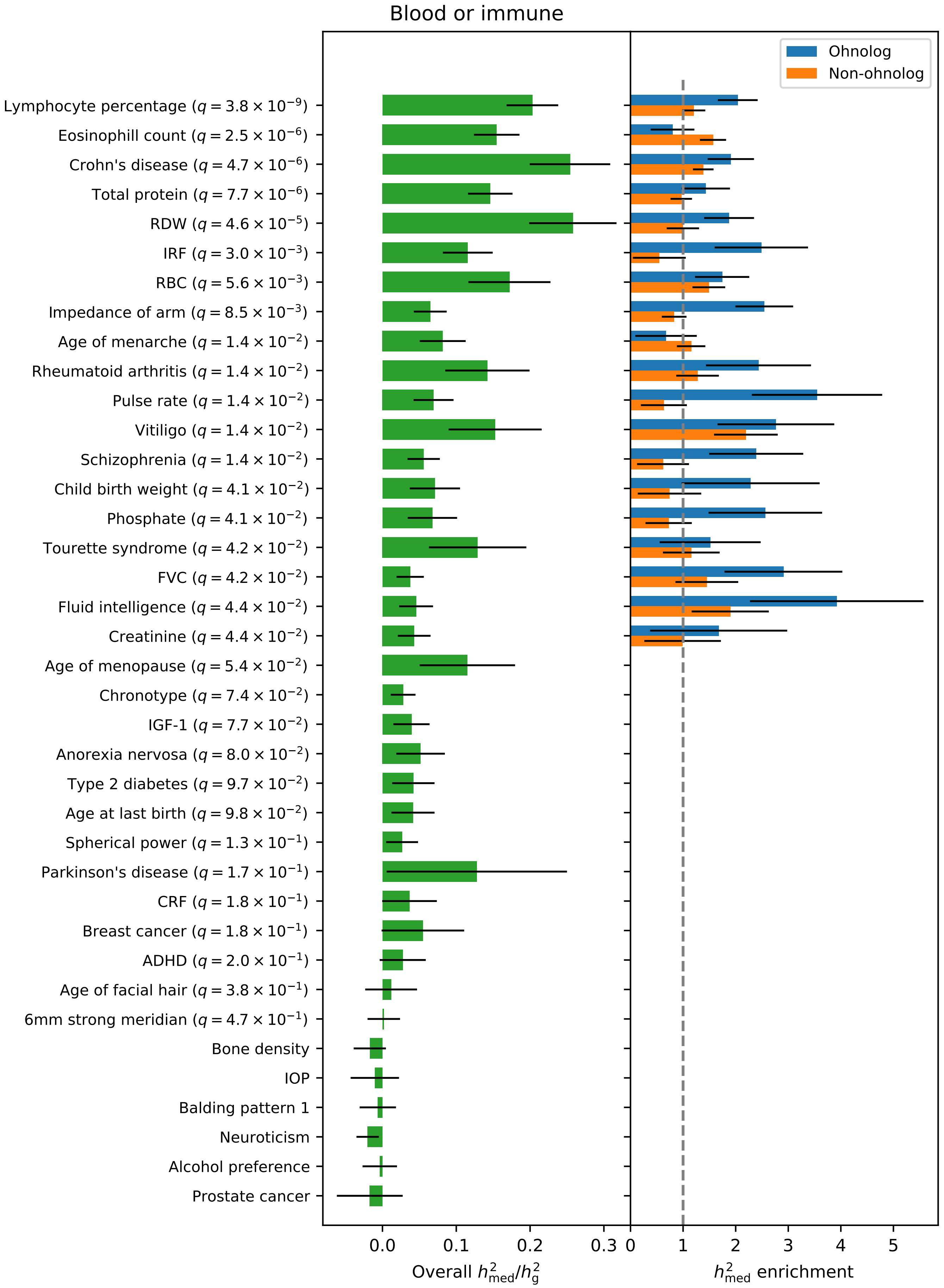

Supplementary Figure 11

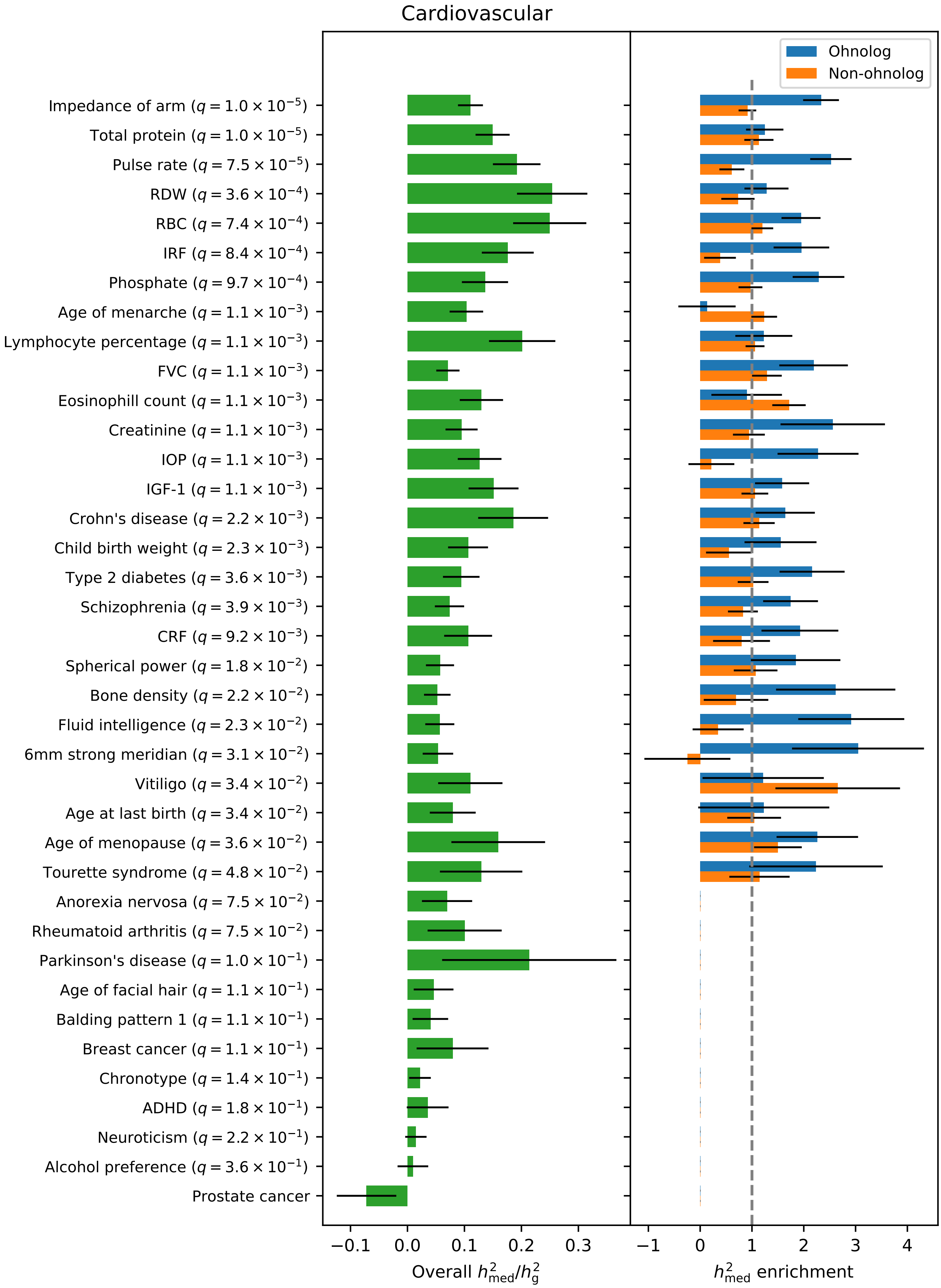

Supplementary Figure 12

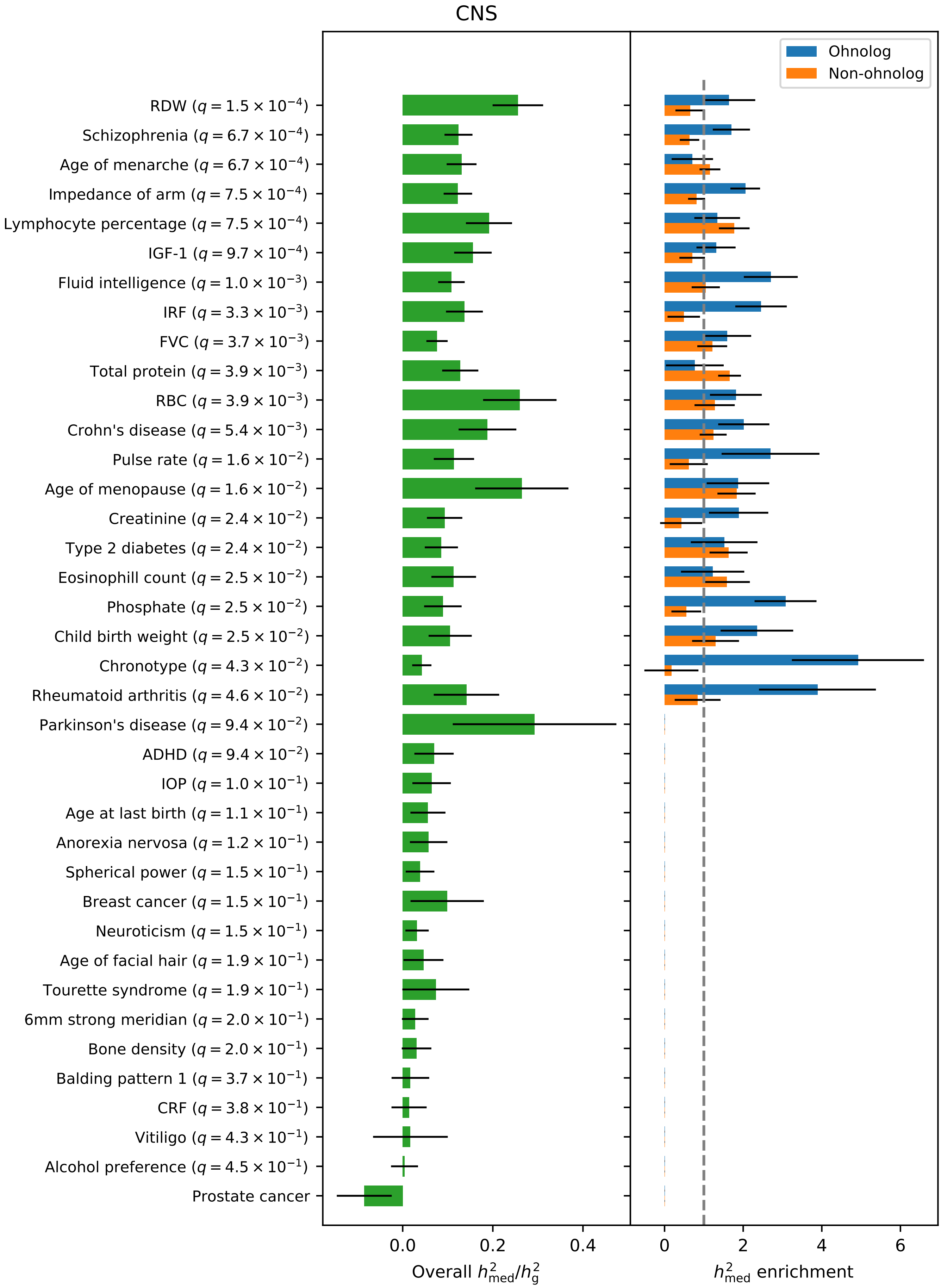

Supplementary Figure 13

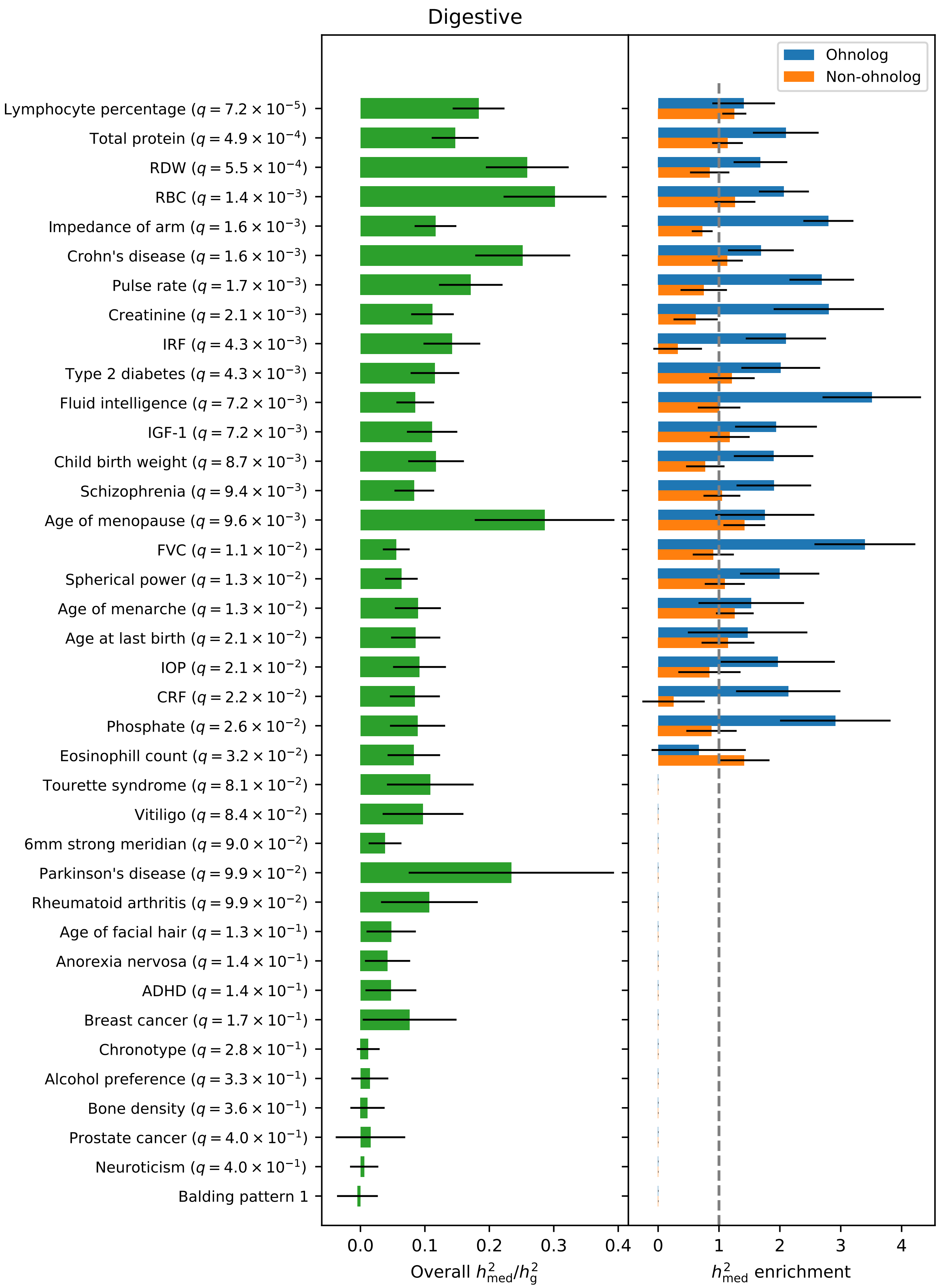

Supplementary Figure 14

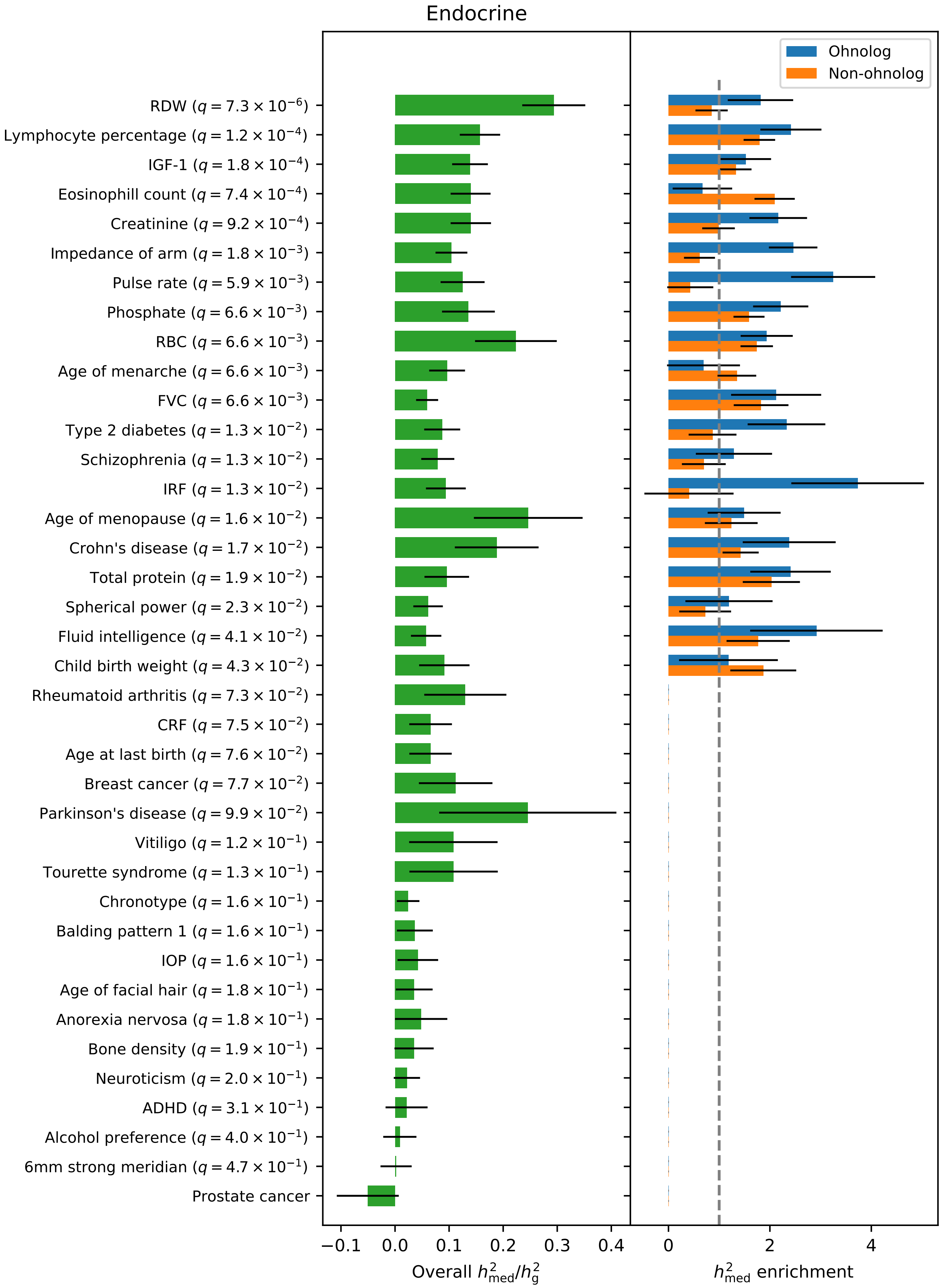

Supplementary Figure 15

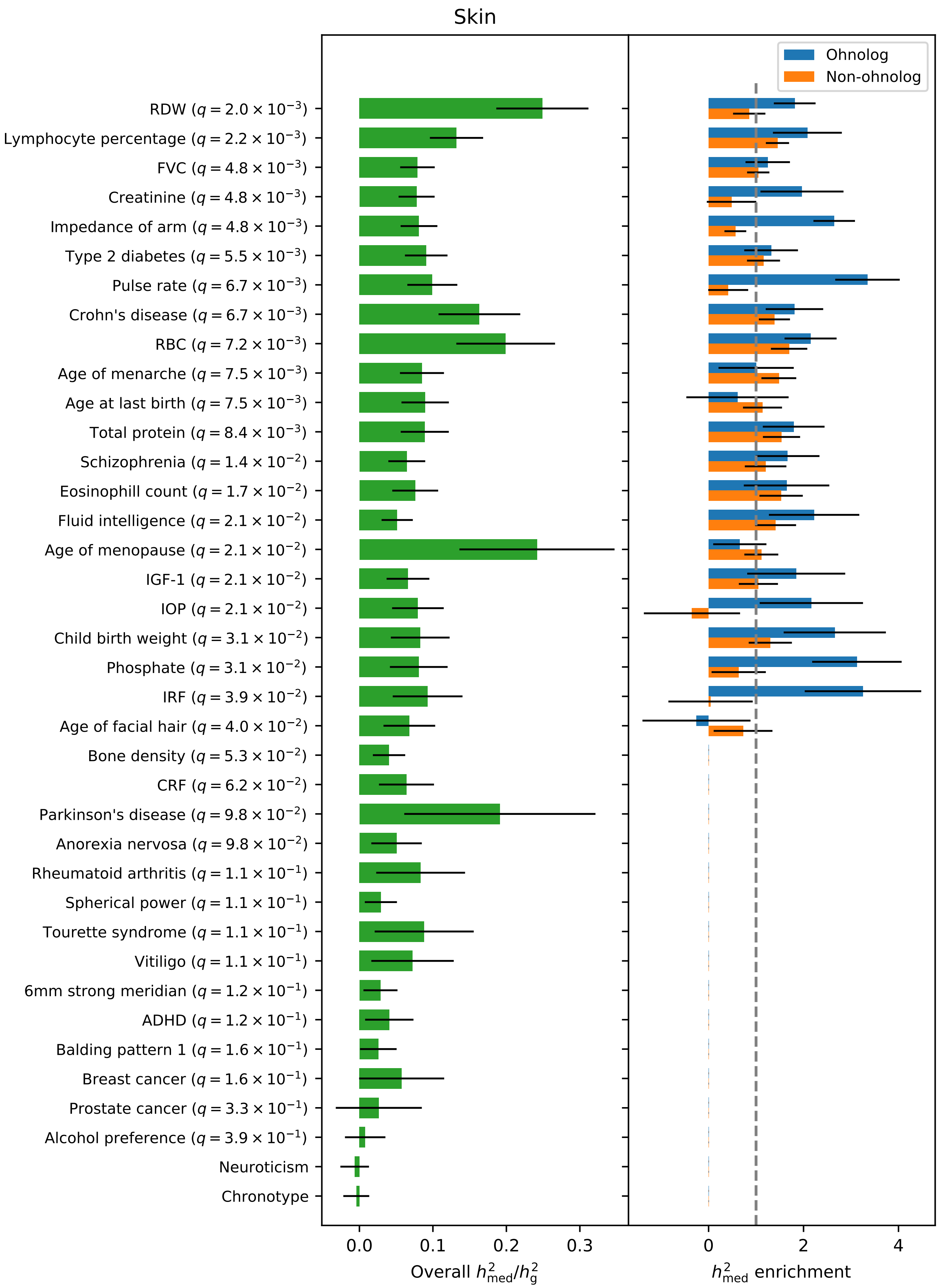
